# Nano-Insecticides Against the Black Cutworm *Agrotis ipsilon* (Lepidoptera: Noctuidae): Toxicity, Development, Enzyme activity, and DNA Mutagenicity

**DOI:** 10.1101/2021.07.23.453522

**Authors:** Mona Awad, El-Desoky S. Ibrahim, Engy I. Osman, Wael H. Elmenofy, Abdel Wahab M. Mahmoud, Mohamed A. M. Atia, Moataz A. M. Moustafa

## Abstract

High-frequency doses of chemical pesticides cause environmental pollution with high pesticide residues. In addition, increasing insecticide resistance in many insect pests requires novel pest control methods. Nanotechnology could be a promising field of modern agriculture, and is receiving considerable attention in the development of novel nano-agrochemicals, such as nanoinsectticides and nanofertilizers. This study assessed the effects of the lethal and sublethal concentrations of chlorantraniliprole, thiocyclam, and their nano-forms on the development, reproductive activity, oxidative stress enzyme activity, and DNA changes at the molecular level of the polyphagous species of black cutworm *Agrotis ipsilon*. The results revealed that *A. ipsilon* larvae were more susceptible to the nano-formsthan the regular forms of both nano chlorine and sulfur within the chlorantraniliprole and thiocyclam insecticides, respectively, with higher toxicities than the regular forms (ca. 3.86, and ca.2.06-fold, respectively). Significant differences in biological parameters, including developmental time and reproductive activity (fecundity and hatchability percent) were also observed. Correspondingly, increases in oxidative stress enzyme activities were observed, as were mutagenic effects on the genomic DNA of *A. ipsilon* after application of the LC_50_ of the nano-forms of both insecticides compared to the control. The positive results obtained here have led us to apply these nano-forms indifferent insect models in additional studies.

## Introduction

The black cutworm, *Agrotis ipsilon* (Lepidoptera: Noctuidae) is a major insect pest that can destroy many important crops worldwide [1]. Specifically, the insect larvae damage many crop species, vegetables, and weeds [2]. *A. ipsilon* larvae can consume over 400 cm^2^ of foliage during their development [3]. Chemical insecticides have been used to prevent crop loss by *A. ipsilon* [4].

The selection of highly effective insecticides and their appropriate application methods is a fundamental problem of integrated pest management strategies for insect pest control. Nanotechnology could provide new methods and agricultural products to counteract the fears related to the potential unwanted environmental impact of chemical insecticides through increased exposure and toxicity in non-target organisms (5). Nanoparticle technology could develop through two distinct mechanisms: (a) supplyingsingularcrop protection or (b) ascarriers for existing pesticides [6]. Generally, nanopesticides are defined as any pesticide formulation of nano-sized, insecticide particles or small, engineered structures with pesticide properties [7,8]. Nanopesticides have several advantages, including increased potency and durability, and a reduced amount of active ingredients [9]. They are also considered to be a promising solution for the reduction of the environmental footprint left by chemical pesticides [7]. The increasing interest in the use of nanopesticides raises questions about their fate, toxicity, and biodegradation [10], as well as how to assess the environmental risk of these materials.

The introduction of nanoinsecticides into the environment necessitates the careful identification of their potential. Generally, insecticide studies focus on evaluating the lethal and sublethal toxicity of insecticides on insect development and enzyme activities [11]. Insecticides are considered a stress factor that may upset the functional balance in insects [known as oxidative stress (OS)]. Insecticides are characterized by the enhanced production of reactive oxygen species (ROS) with the simultaneous impairment of their scavenging systems. Increased ROS concentrations result in oxidative damage to proteins, lipids, and nucleic acids, and thus cell function, organs, and the entire organism may be seriously disrupted, resulting in death [12]. To avoid, or at least reduce this effect, organisms have developed effective defense systems controlled by their nervous and endocrine glands. The oxidative stress enzymes are an essential group of enzymes that combat the adverse effects of ROS on cells. Several defense mechanisms have been developed by insects and include enzymatic and non-enzymatic components [13,14]. Superoxide dismutase (SOD), catalase (CAT), peroxidase (POX), glutathione (L-ɣ-glutamyl-L-cysteinylglycine, GSH) are major antioxidant enzymes in insects that play a fundamentalrole in cell protection by removing oxidative stress [15]. SOD is an antioxidant enzyme, which converts superoxide into oxygen and hydrogen peroxide [16]. CAT is mainly aH_2_O_2_-scavenging enzyme that principally removes H_2_O_2_ generated from developmental or environmental stimuli into water and oxygen in all aerobic organisms [17]. CAT tends to reduce small peroxides, such as H_2_O_2_, but does not affect larger molecules, such as lipid hydroperoxides. POX utilizes either H_2_O_2_or O_2_to oxidize a wide variety of molecules [18], and uses H_2_O_2_ to oxidize phenolic compounds. GSH is the heart of the essential cellular antioxidant system [19], and may serve as an electron donor (cofactor) for antioxidant enzymes like glutathione peroxidases and glutathione S-transferases [20].

Genetic molecular markers have become a central tool to determine the degree of genetic variability [21], molecular phylogenetics [22], genetic fidelity [23], and disease resistance [24] in organisms. Various kinds of molecular marker techniques have been identified in insect populations [25]. The inter-simple sequence repeats (ISSR) are a useful marker tool to detect genetic variation and differentiate closely-related individuals [26]. The high variability level of the ISSR marker has been indicated to be a common characteristic of Lepidoptera genomes and is applied as a fingerprint technique between groups of organisms [27].

Due to the lack of information on nano-insecticide toxicity, several studies are required to better understand their effects on the biological and physiological parameters of target insects [10]. There are currently no sufficient screening methods to assess whether nanoinsecticides are safe for field administration without significant side effects on human health. Thus, this study’s main goal was to evaluate the potential effects of the nano-forms of two new insecticides, chlorantraniliprole and thiocyclam, on *A. ipsilon*. We also investigated the effects of the lethal and sublethal concentrations of both insecticides and their nano-forms on the development, reproductive activity, and oxidative stress enzyme activities, including; SOD, CAT, lipid peroxidase, and GR of *A. ipsilon*. This study examined both insecticides for nanoparticle-induced changes in *A. ipsilon*at the DNA molecular level.

## Materials and Methods

### Insect rearing

*Agrotis ipsilon* was reared in a rearing room at 26 ± 1 °C, 65 ± 5% relative humidity under a reversed 16 L:8 D. The newly hatched larvae were kept in a clean glass jar (1 L) and provided castor oil leaves daily until the third instar larvae emerged. They were then transferred to larger, clean glass jars (2 L) to prevent larval cannibalism. The bottom of each jar was covered with a thick layer of fine sawdust and the usual rearing techniques were performed along with the developing instars larvae till pupation occurred. The emerged moths were supplied with a 10% sugar solution as supplement dietary [11].

### Insecticides and chemicals

Chlorantraniliprole (Coragen^®^ 20% SC, suspension concentrate, DuPont) and thiocyclam (Evisect-S^®^ 75% SC) were tested. All chemicals used in the preparation of the nano-form of the insecticides were purchased from Sigma Chemical Co. (St. Louis, USA) without further purification. The oxidative stress enzymes kits were purchased from Biodiagnostic Company, Egypt.

### Nano-insecticides preparation and size measurements

Nano-chlorine (chlorantraniliprole) and nano-sulfur (thiocyclam) were prepared by [28,29,30], with some modifications. Hydrochloric acid (HCl) and orthorhombic bravais were used as a chlorine source, while sodium thiosulphate and sulfuric acid were used as a sulfur source. The nano-chlorine was made from a sedimentarysolution of HCl and MnO2 v:v, where hydrochloric acid and manganese oxide were added slowly in a molar ratio (3:2) in the presence of the stabilizing agent PVA using forceful moving for 5 hours. The obtained precipitation was filtered and washed thoroughly with deionized water, then the suspension solution was added to 16 ml of NaCl 0.2 M aqueous solution using forceful moving at a steady rate for40 min. The reaction fusion was stirred for an additional 3 hours at ambient temperature. The precipitate was mixed with orthorhombic brava is w:w (2:1) in the presence of HCl 90%, then centrifuged at 1500 rpm for 30 min. Next, the solution was cooled in an ice bath, and then subsequently exposed to 1.5 psi pressure continuously for 6 hours.

Nano-sulfur was prepared from a sedimentary solution of sodium thiosulphate and sulfuric acid (1:1). Sodium polysulfide and hydrochloric acid solutions were incrementally mixed in a molar ratio of 3:2 under forceful moving for 8 hours. The obtained precipitation was filtered and washed methodically with deionized water in a mixed water/toluene system, then washed with ionized water for 3 hours. The precipitation was mixed with oxalic acid 1 M and trimethylammoniumbromide compound solution (molar ratio 1:3) under slow stirring at 33 °C for 6 hours. Afterwards, drop benzene sulphonate (molar ratio 2:3) was added, and the resulting solution was kept at 1.5 psi pressure for 3 days discontinuously (7 hours per day). Finally, the solution was dried in an oven at 90 °C for continuously for 3 days. The final nano-suspension was prepared in deionized water and left on a shaker for 2 days at 20 °C.

### Bioassays

The toxicity of chlorantraniliprole, thiocyclam, and their nano-forms on the second instar larvae were assessed using the leaf dipping technique [31], with some modifications. Briefly, six different concentrations were prepared for each, and water was used to control the larvae. Treated and untreated castor bean leaves were transferred after drying into a glass jar (0.25 L), and 10 larvae were added to each jar with five replications and left to feed for 24 h. Afterwards, all larvae were offered untreated leaves, and the mortality% was recorded four days (96 hours) post treatment [32] to calculate the lethal and sublethal concentrations for each insecticide form. The bioassay was repeated twice.

### Effects on *A. ipsilon* development

The sublethal and lethal concentrations (LC_15_ and LC_50_) for each form of chlorantraniliprole and thiocyclam were used on the second instar larvae using the method described above. Surviving insects were used to study the effect of each insecticide on larval and pupal developmental time, pupation%, and adult emergence. Seven days after exposure, the surviving larvae were transferred individually to a clean cup to record the development time of the larval and pupal stages and the pupation%. After pupation, each pupa was sexed, weighed, and kept individually in the same cup to record the emergence%.

### Studies on fecundity and fertility

Groups of 5 females and 7males in 3 replicates [33] were used to calculate the number of eggs and hatching% after the second instar larvae were treated with the LC_15_ and LC_50_ values of each insecticide and their nano-forms. Deposited eggs were collected and counted on days 2–6 in the mating jars. The eggs were transferred to a clean jar and kept for 5 days to record the hatching%.

### Oxidative stress enzyme assays

#### Sample preparation

Seven days after LC_15_ and LC_50_ equivalent treatment of the second instar larvae, 100 mg fresh body weight of the surviving larvae were transferred to clean and sterilize Eppendorf tubes (1.5 ml). The samples were stored immediately at −20 °C until later analysis. Each treatment and control was replicated five times. The treated larvae were homogenized in a potassium phosphate buffer (50 mM, pH 7.0) at30 µl buffer per 1 mg of body weight. The homogenate was centrifuged for 15 min at 7000 g at 4 °C, and the supernatants were used for further analysis.

#### Enzymes measurement

SOD activity was determined [34] at the absorbance of 560 nm. The CAT enzyme activity was estimated by measuring the rate of H_2_O_2_ consumption [35] via absorbance at 510 nm. The level of lipid peroxidase was assayed by monitoring the formation of malondialdehyde (MDA) at 534 nm [36]. GR activity was estimated as the reduced glutathione (GSSG) in the presence of NADPH [37], which oxidizes to NADPH^+^ at 340 nm. The total protein concentration of all samples was measured spectrophotometrically based on the Biuret Method using Protein Biuret Kit (Biodiagnostic, Egypt).

### Molecular analysis

#### DNA extraction

DNA was isolated from both treated and untreated second instar larvae using a G-spin™ total DNA extraction kit (INtRON Biotechnology) per the manufacturer’s instructions. The DNA was quantified with a Qubit 4 Fluorometer (Thermo Fisher Scientific Inc.). The DNA concentrations were measured and subsequently adjusted in all samples to 10 ng/µL for subsequent molecular analyses.

#### ISSR polymorphism analysis

For ISSR PCR amplification [38,39], a set of 15 ISSR primers were applied against the 9 treatments. The PCR was carried out in a total volume of 25 μl containing the following components: 25 ng genomic DNA; 1X PCR buffer; 1.5 mM MgCl_2_; 0.25 mM of each dNTPs; 1 μM of each primer; 1 U Go-Taq Flexi polymerase (Promega).

Thermocycling amplification was performed with a GeneAmp PCR system 9700 (Applied Biosystem, Inc.). The amplification was programmed at 94 °C for 5 min for the initial denaturation cycle, followed by 35 cycles with each cycle comprised of (94 °C for 1 min, 50 °C for 1 min, then 72 °C for 90 s) and a final extension at 72 °C for 7 min. The produced PCR amplicons were electrophoresed using 1.5% agarose gel. A 100 bp plus DNA ladder and 1 kb were used as molecular size standards. PCR products were photographed using a Gel Doc™ XR+ System (Bio-Rad®).

### Data analysis

#### Biological and biochemical data analyses

The statistical analysis program LDP line was used to determine the lethal and sublethal concentration values (LC_15_, LC_50_, and LC_90_) for each insecticide and its nano-forms (with 95% confidence limits). All biological parameters and oxidative stress enzymes activity (SOD, CAT, lipid peroxidase, and GR) were performed using one way ANOVA in addition to Dunnett’s multiple comparisons test with Graph Pad Prism 8 statistical analysis software. Moreover, a silhouette analysis was performed to evaluate the quality of the reproductive activity, enzymes, and developmental measurements by testing the cluster distances within and between each cluster [40]. Additionally, we performed a multidimensional preference analysis to disclose the interrelationships among parameters in addition to the similarity classification in terms of dependent and independent variables in different space dimensions [41]. Finally, hierarchical clustering based on the correlation analysis was conducted with two-dimensional heatmap plotting was constructed.

#### Molecular data analysis

For ISSR data analysis, the generated amplicons were scored visually. To generate a binary data set, the amplicons were scored as absent (0) and present (1). The polymorphism percentage was analyzed by dividing the number of amplified polymorphic bands by the total number of amplified bands separately for each primer [42]. A similarity matrix was built to estimate the genetic distances between all possible treatment pairs. The Jaccard coefficient was used for the pairwise comparisons [43]. The genetic similarity (GS) between each pair of treatments was calculated using GS = a/(n-d), in which n is the total number of fragments; a is the number of positive coincidences; and d is the number of negative coincidences. The genetic distances (GD) between pairs of treatments were estimated using GD = 1-GS. The unweighted pair group method of arithmetic averages (UPGMA) was used to construct the dendrogram [44].

The efficiency of the ISSR primers was determined by calculating the following parameters: expected heterozygosity (H = 1 – Σ pi^2^ according to [45]; polymorphism information content (PIC = 1 – Σ pi^2^ – Σ pi^2^) according to [46]; effective multiplex ratio (E = n β) according to [47]; marker index (MI = E Hav) according to [47]; mean heterozygosity (Hav = Σ H_n_/n_p_) according to [47]; discriminating power (D = 1 – C) according to [48]; resolving power (R = Σ I_b_) according to [49].

## RESULTS

### Characterization of nano-(chlorine) chlorantraniliprole and nano-(sulfur) thiocyclam

The nano-suspensions of chlorine (Fig 1) and sulfur (Fig 2) were achieved using top-down molecular chemical techniques. Briefly, a single drop of the nanoparticle solution was spread onto a carbon-coated copper grid, and then was posteriorly dried at room temperature for transmission electron microscope (TEM) analysis. The dimensions of the nanoparticles were established directly from the figure using Image-Pro Plus 4.5 software. The particles were irregular in shape, with dimensions of 3.99 nm for chlorine and 4.05 nm for sulfur (Table 1), as well as 98.5% purity for each element. The nanoparticles’ shape and dimension were examined using a JEOL 1010 TEM at 80 kV (JEOL, Japan).

**Table 1.** Particle size of nano-chlorine, and nano-sulphur

| Formulation | Sample number | Size (nm) | Average size (nm) |
| --- | --- | --- | --- |
| Nano-chlorine | 1 | 3.25 | 3.99 |
|  | 2 | 7.11 |  |
|  | 3 | 3.17 |  |
|  | 4 | 2.46 |  |
| Nano-sulphur | 1 | 3.84 | 4.05 |
|  | 2 | 4.41 |  |
|  | 3 | 3.73 |  |
|  | 4 | 4.22 |  |

**Fig 1.**
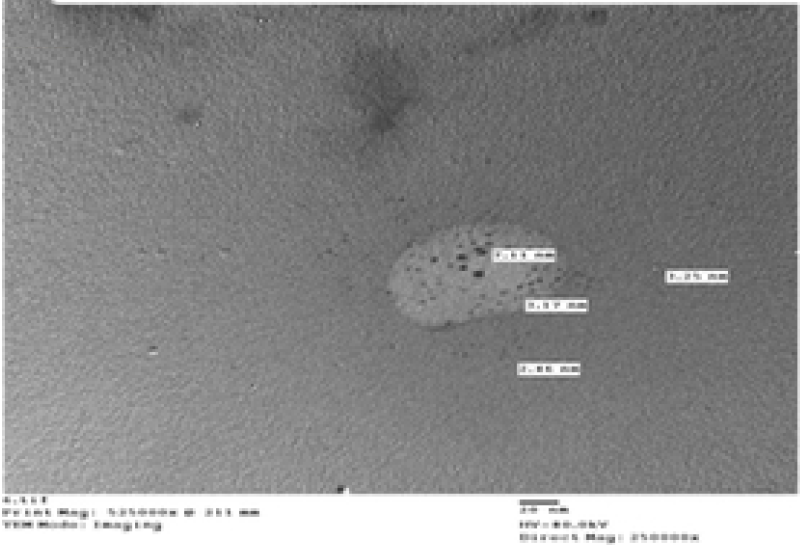
Nano-chlorine

**Fig 2.**
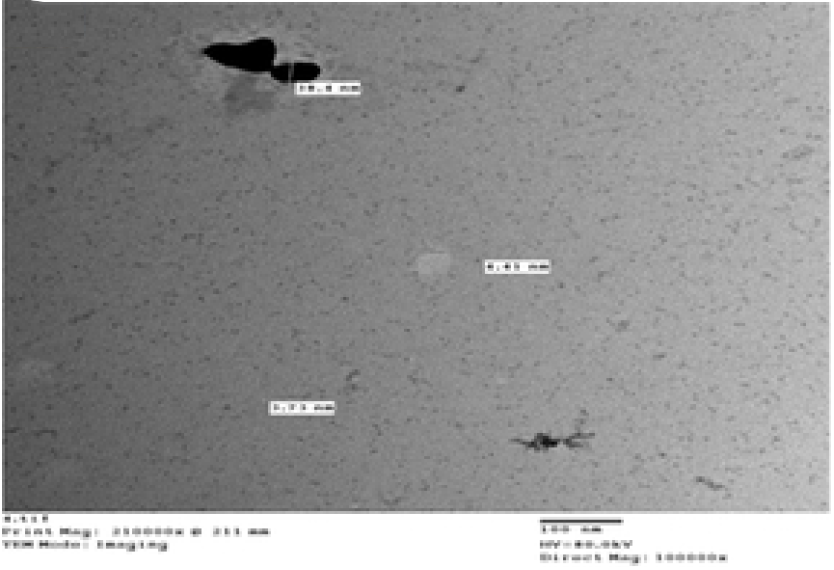
Nano-sulphur

### Activities of chlorantraniliprole, thiocyclam, and their nano-forms

The toxicity of chlorantraniliprole, thiocyclam, and their nano-forms on the second instar larvae are presented in Table 2. The LC_50_ values were 0.058 and 9.20 mg/l for chlorantraniliprole and thiocyclam, respectively, 96 h post treatment. In contrast, the nano-forms (nano-chlorantraniliprole and nano-thiocyclam) had significantly higher toxicities than their original compounds, with LC_50_ values of 0.015 and 4.46 mg/l, respectively (Table 2).

**Table 2.**
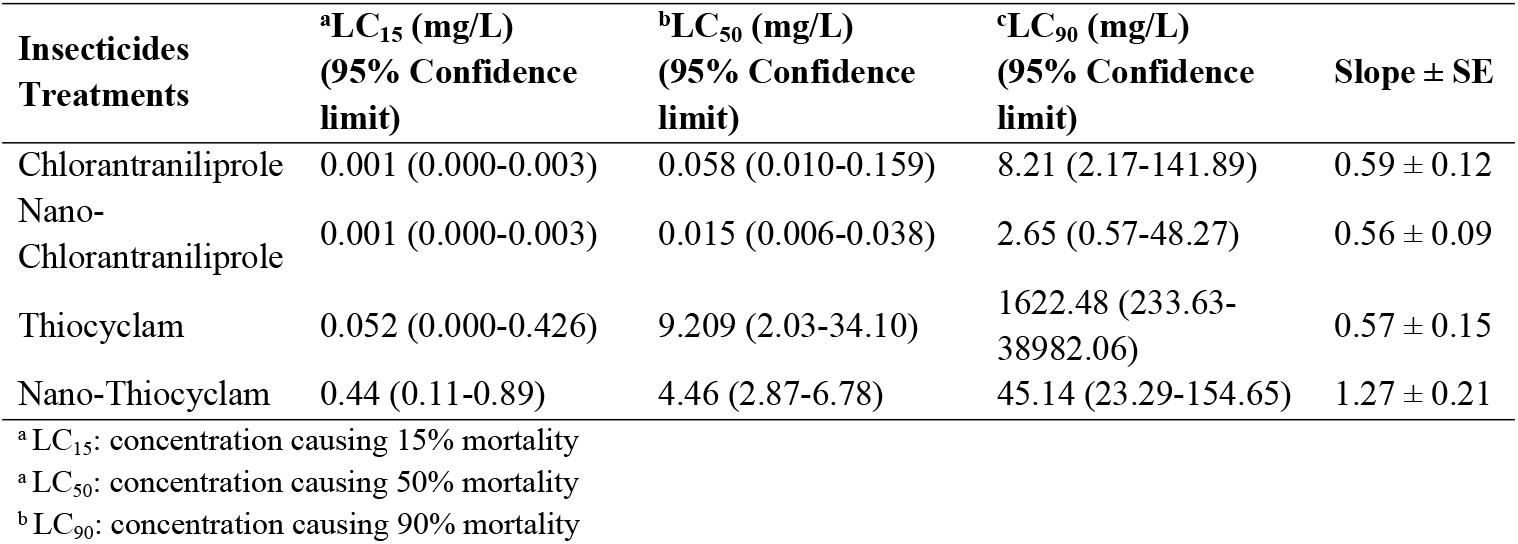
Lethal and sublethal activity of chlorantraniliprole, thiocyclam, and their nano-form in the 2^nd^ instar larvae of *Agrotis ipsilon*

### Lethal and sublethal effects on the second instar larvae

Table 3 shows the latent effect of chlorantraniliprole, thiocyclam, and their nano-forms on *A. ipsilon* development (from second instar larvae till the emergence) due to LC_15_ and LC_50_ exposure. The results showed elongation of the larvae developmental period under all LC_50_treatments, but only a slight significance in the pupal stage duration. The pupation% decreased significantly after LC_15_ and LC_50_ treatmentof nano-thiocyclam at 83.20 ± 4.36% and 85.88 ± 0.56%, respectively (Table 3). Low significant differences were found in the female pupal weight under all treatments. No differences were found in the male pupal weight, sex ratio, or emergence% under all treatments (Table 3).

**Table 3.** Effects of chlorantraniliprole, thiocyclam, and their nano-form in the developmental stages of *A. ipsilon*.

| Treatments |  | *Larval duration | Pupation% | **Pupal duration | Pupal weight (mg) |  | Sex ratio |  | Emergence % |
| --- | --- | --- | --- | --- | --- | --- | --- | --- | --- |
|  |  |  |  |  | Female | Male | Female | Male |  |
| Control |  | 20.22±0.15 | 93.79±3.10 | 17.00±0.15 | 0.43±0.01 | 0.39±0.01 | 43.44±3.79 | 56.56±3.79 | 98.61±1.39 |
| Chlorantraniliprole | LC<sub>15</sub> | 21.04<sup>ns</sup>±0.29 | 95.84<sup>ns</sup>±0.31 | 17.91<sup>**</sup>±0.16 | 0.40<sup>ns</sup>±0.01 | 0.39<sup>ns</sup>±0.01 | 49.55<sup>ns</sup>±7.58 | 50.45<sup>ns</sup>±7.58 | 98.61<sup>ns</sup>±1.39 |
|  | LC<sub>50</sub> | 24.13<sup>****</sup>±0.46 | 90.16<sup>ns</sup>±0.15 | 18.48<sup>****</sup>±0.28 | 0.39<sup>ns</sup>±0.01 | 0.35<sup>ns</sup>±0.01 | 42.85<sup>ns</sup>±2.28 | 57.15<sup>ns</sup>±2.28 | 98.15<sup>ns</sup>±1.85 |
| Nano-Chlorantraniliprole | LC<sub>15</sub> | 22.25<sup>***</sup>±0.31 | 93.92<sup>ns</sup>±1.31 | 17.73<sup>ns</sup>±0.24 | 0.39<sup>ns</sup>±0.01 | 0.36<sup>ns</sup>±0.01 | 49.38<sup>ns</sup>±9.97 | 50.62<sup>ns</sup>±9.97 | 96.74<sup>ns</sup>±1.63 |
|  | LC<sub>50</sub> | 22.30<sup>****</sup>±0.26 | 91.07<sup>ns</sup>±2.25 | 17.46<sup>ns</sup>±0.20 | 0.42<sup>ns</sup>±0.01 | 0.40<sup>ns</sup>±0.01 | 51.78<sup>ns</sup>±10.26 | 48.22<sup>ns</sup>±10.25 | 90.54<sup>ns</sup>±2.32 |
| Thiocyclam | LC<sub>15</sub> | 21.66<sup>*</sup>±0.34 | 92.60<sup>ns</sup>±1.23 | 17.77<sup>*</sup>±0.19 | 0.39<sup>ns</sup>±0.02 | 0.38<sup>ns</sup>±0.01 | 42.15<sup>ns</sup>±5.32 | 57.85<sup>ns</sup>±5.32 | 98.24<sup>ns</sup>±1.75 |
|  | LC<sub>50</sub> | 23.30<sup>****</sup>±0.39 | 93.82<sup>ns</sup>±1.91 | 17.46<sup>*</sup>±0.16 | 0.41<sup>ns</sup>±0.01 | 0.37<sup>ns</sup>±0.01 | 46.40<sup>ns</sup>±1.32 | 53.60<sup>ns</sup>±1.32 | 98.61<sup>ns</sup>±1.39 |
| Nano-Thiocyclam | LC<sub>15</sub> | 22.52<sup>****</sup>±0.28 | 86.54<sup>ns</sup>±1.73 | 17.54<sup>ns</sup>±0.19 | 0.36<sup>*</sup>±0.008 | 0.35<sup>ns</sup>±0.01 | 42.67<sup>ns</sup>±4.66 | 57.33<sup>ns</sup>±4.66 | 96.63<sup>ns</sup>±1.71 |
|  | LC<sub>50</sub> | 22.75<sup>****</sup>±0.35 | 82.55<sup>*</sup>±3.79 | 17.27<sup>ns</sup>±0.12 | 0.39<sup>ns</sup>±0.01 | 0.39<sup>ns</sup>±0.01 | 49.59<sup>ns</sup>±12.14 | 50.41<sup>ns</sup>±12.14 | 88.85<sup>*</sup>±3.38 |

Values marked with the same letters are not significantly different ( $P > 0.05$ : Dunnett's multiple comparisons test).
\* number of days from 2<sup>nd</sup> instar larvae till pupation
\*\* number of days from the pupation till the emergence

### Fecundity and fertility

Chlorantraniliprole, thiocyclam, and their nano-forms significantly decreased the hatchability percent under LC_15_ and LC_50_compared to the control (Table 4). After chlorantraniliprole and nano-chlorantraniliprole treatment, the hatchability percentages were 84.95, and 81.29% at LC_15_, and 73.83, and 77.74% at LC_50,_ respectively. Under thiocyclam and nano-thiocyclam treatment, the hatchability percentages were 82.64, and 78.32% at LC_15_, and 78.79, and 71.99% at LC_50_. In contrast, the number of eggs laid by one female (fecundity) showed no significant differences between treated and untreated larvae (Table 4).

**Table 4.**
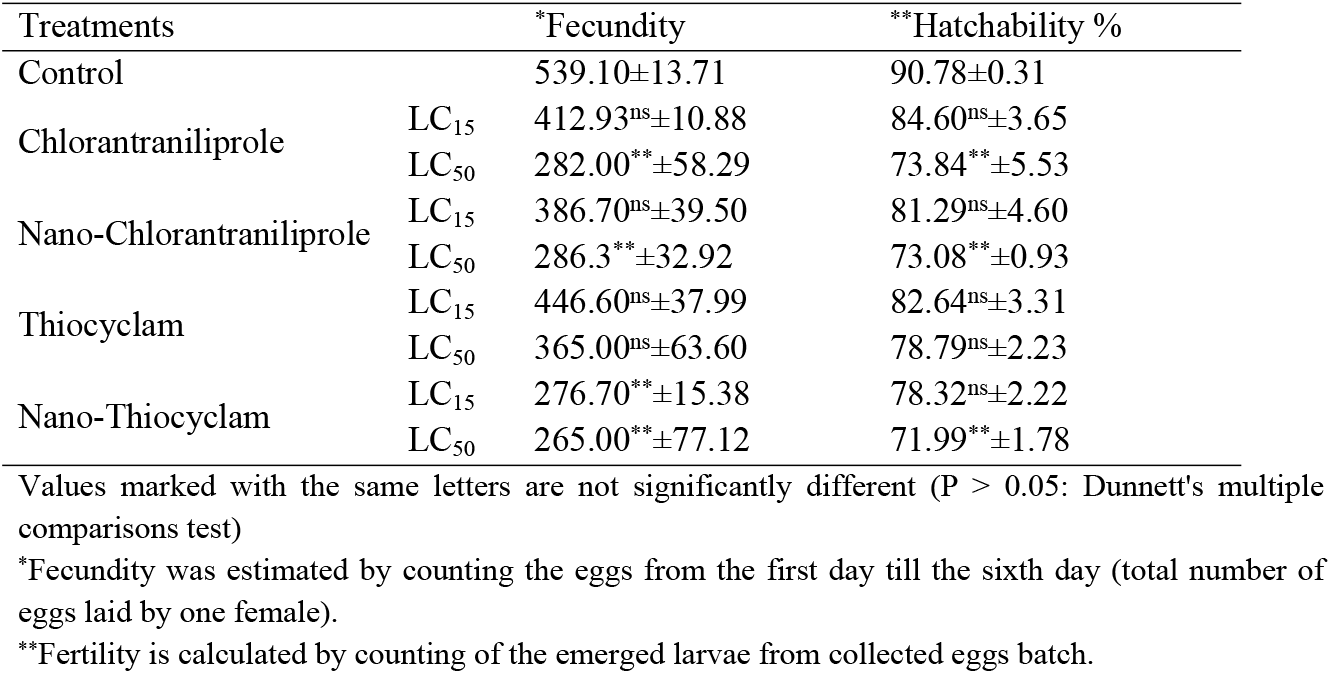
Mean fecundity and hatchability % (±SE) of *A. ipsilon* females after treated the 2^nd^ instar larvae with LC_15_ and LC_50_ values of chlorantraniliprole, thiocyclam, and their nano-form.

### Activity of oxidative stress enzymes

Table 5 shows that exposure to both insecticides and their nano-forms caused a significant increase in SOD activity after treatment with the LC_50_ of thiocyclam (40.09 U/g of protein) and nano-thiocyclam (43.0 U/g of protein). The SOD activity after treatment with the LC_50_ of chlorantraniliprole, and nano-chlorantraniliprole were highly significant compared to the control treatment with 45.38 and 53.96 U/g of protein (ca. 5.44, and 6.47 fold), respectively. No significant changes relative to the control were recorded in any other treatments.

**Table 5.**
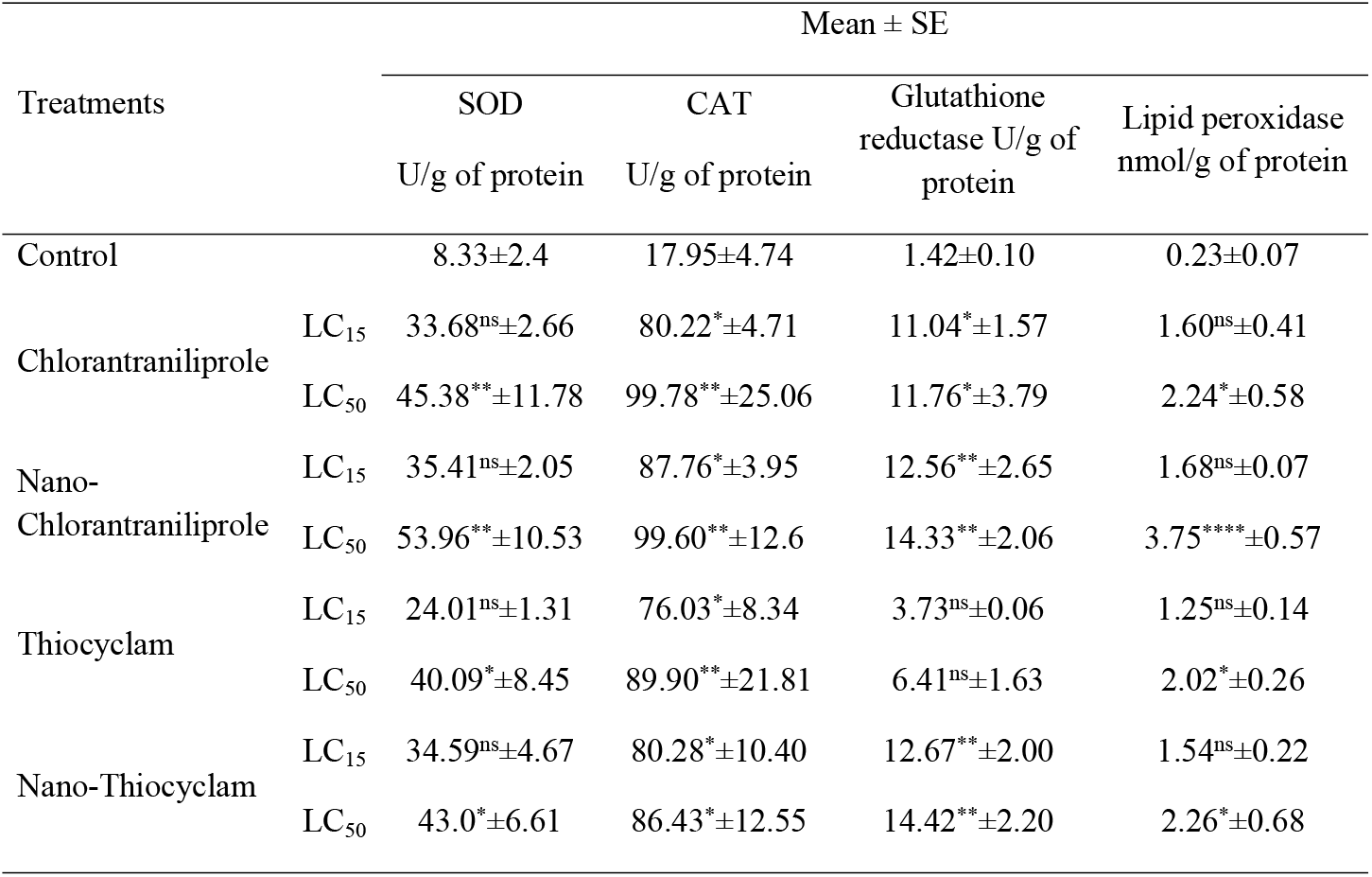
Mean (±SE) of oxidative stress enzymes (SOD, CAT, glutathione reductase, and lipid peroxidase) activities of *A. ipsilon* after exposure of 2^nd^ instar larvae to LC_15_ and LC_50_ values of chlorantraniliprole, thiocyclam, and their nano-forms.

The LC_15_ and LC_50_ of both insecticides and their nano-forms stimulated CAT activity compared to the control (Table 5). A significant increase in CAT activity was observed at the LC_15_ of chlorantraniliprole and nano-chlorantraniliprole, the LC_15_ of thiocyclam, and both theLC_15,_ and LC_50_ of nano-thiocyclam (80.22, 87.76, 76.03, 80.28 and 86.43 U/mg of protein, respectively). The highest significance was observed for the LC_50_ of chlorantraniliprole and nano-chlorantraniliprole, and the LC_50_ of thiocyclam (ca. 5.56, 5.54, and 5.0-fold, respectively).

An increase in lipid peroxidase activity was observed under exposure totheLC_50_ of nano-chlorantraniliprole with 3.75 nmol/g of protein (ca. 16.30-fold). Meanwhile, the lipid peroxidase activity after treatment with the LC_50_ of chlorantraniliprole, thiocyclam, and nano-thiocyclam were slightly significantly at 2.24, 2.02, and 2.26 nmol/g of protein, respectively. No significant differences in lipid peroxidase activities were observed in any other treatments (Table 5).

Table 5 shows that exposure to all the investigated insecticides and their nano-forms resulted inthe significant stimulation of GR activity in the LC_15_ and LC_50_ of chlorantraniliprole (11.4 and 11.76 U/g of protein). GR activity was significantly high under the LC_15_ and LC_50_ nano-chlorantraniliprole treatments at 8.84 and 10.09-fold, respectively and under the LC_15_ and LC_50_ nano-thiocyclam treatments at 8.92, and 10.15-fold, respectively. No significantchangesin enzyme activity was observed for the LC_15_ and LC_50_thiocyclam treatments (3.73 and 6.41 U/g of protein) compared to the control.

### Correlation between development, reproductive, and enzyme activity

The plots for silhouette analysis were calculated based on the Euclidean distance metric to assess the cluster quality of the treatments based on reproductive activity, enzymes, and developmental measurements via cluster distance tests within and between each cluster (Fig 3). The results revealed that all parameters, except GR, SOD, and CAT enzymes, exhibited negative values, thus indicating that the clusters were mostly similar, and that the cluster configuration may have too few clusters. Meanwhile, two-dimensional heatmap plotting based on all parameters clustered the LC_15_ of thiocyclam, chlorantraniliprole, and nano-chlorantraniliprole as the most similar to the control (1^st^ cluster), whereas the LC_50_ of nano-chlorantraniliprole, nano-thiocyclam, chlorantraniliprole, and the LC_15_ of nano-thiocyclam were grouped in the second cluster as less-similar relative to the control (Fig 4). Moreover, multidimensional preference analysis was performed to summarize the inter relationships of all treatments, parameters, and classes (Fig 5). The plot shows that the LC_50_ of nano-thiocyclam, chlorantraniliprole, and the LC_15_ of nano-thiocyclam deviated the most compared to the control. GR, SOD, and CAT enzymes revealed almost the same pattern.

**Fig 3.**
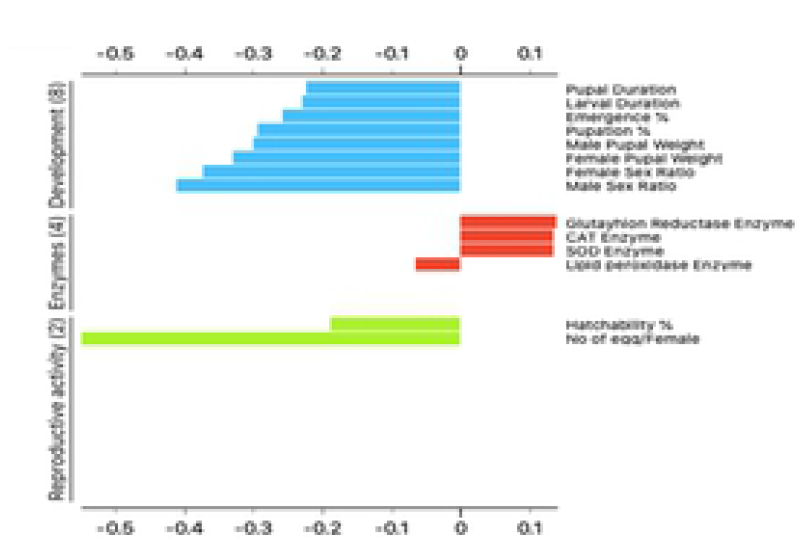
Plot of Silhouette analysis values for clustering of Reproductive activity, Enzymes and Developmental variables. On the y-axis each cluster are ordered by decreasing silhouette value. The silhouette value can range between −1 and 1.

**Fig 4.**
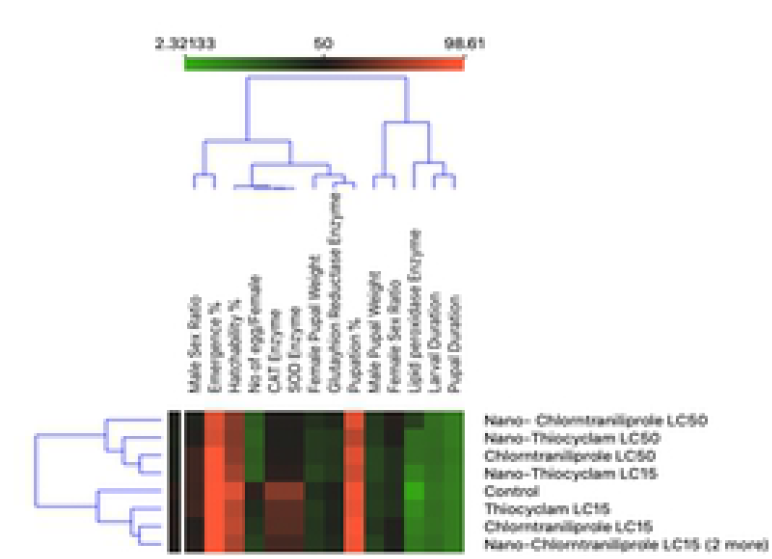
Two-dimensional heatmap visualization shows the interaction between the treatments and (A) the eight developmental parameters (B) the two reproductive activity parameters (C) the four enzymes parameters.

**Fig 5.**
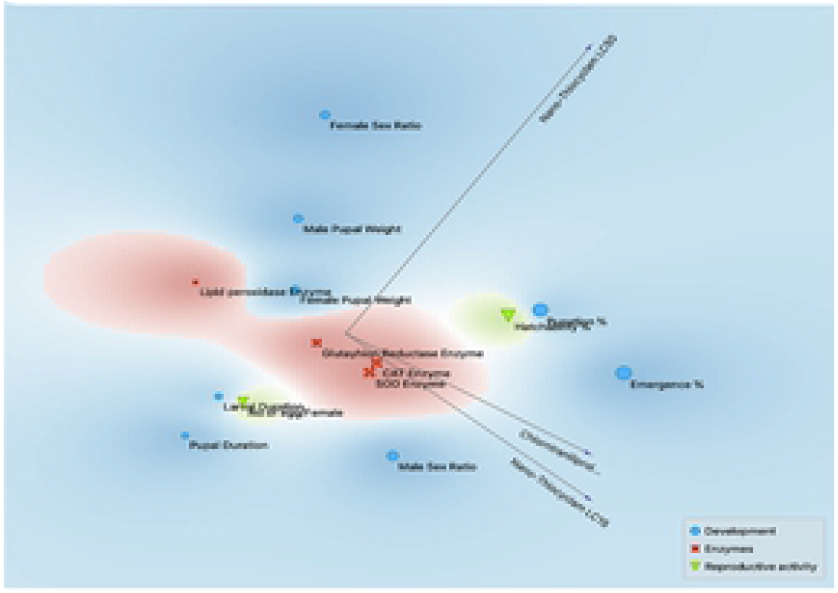
Multidimensional preference analysis plot summarizing the interrelationships amongst treatments, parameters, and classes.

### ISSR analysis

To determine the mutagenic levels of the chlorantraniliprole, thiocyclam, and their nano-forms on the DNA of the second instar larvae, the ISSR marker system was used. The DNA of the untreated second instar larvae and all other treatments were amplified using 15ISSR primers (Fig 6). The 15 ISSR primers yielded a total of 252 scorable amplicons with an average of 16.8 bands/primer (Table 6). The number of amplified DNA fragments per primer ranged from 11 bands (primer ISSR-18) to 21 bands (primers ISSR-10 and ISSR-12). The number of polymorphic bands per primer ranged from 7 bands (primer ISSR-1) to 19 bands (primer ISSR-5). A narrow range of the expected heterozygosity values was observed between 0.37 to 0.50. Notably, 13 out of the 15 ISSR primers showed values near 0.50. The polymorphism information content almost revealed the same pattern for all primers, with values ranging from 0.30 to 0.37. The effective multiplex ratio values were almost all high, with values ranging from 6.00 to 12.67. In contrast, the marker index values were shallow (near to 0.01). The discriminating power values ranged from 0.44 to 0.86. The resolving power values ranged between 4 to 10. The primers ISSR-2 and ISSR-5 showed the highest-resolving power value (10.44) among all the ISSR primers (Table 6). The highest GS was observed between the LC_50_ of chlorantraniliprole and the LC_15_ of thiocyclam. The lowest GS was determined to be between the LC_50_ of nano-chlorantraniliprole, and nano-thiocyclam (Table 7).

**Table 6.** Primer names, number of total bands, polymorphic bands, percentage of polymorphism and markers efficiency parameters of ISSR primers.

| Primer Name | No. of Polymorphic Bands | No. of Monomorphic Bands | Total No. of bands | % of polymorphism | H | PIC | E | H.av | MI | D | R |
| --- | --- | --- | --- | --- | --- | --- | --- | --- | --- | --- | --- |
| ISSR-1 | 7 | 6 | 13 | 53.8 | 0.37 | 0.30 | 9.78 | 0.00 | 0.03 | 0.44 | 4.00 |
| ISSR-2 | 15 | 2 | 17 | 88.2 | 0.50 | 0.37 | 8.56 | 0.00 | 0.03 | 0.75 | 10.44 |
| ISSR-3 | 18 | 1 | 19 | 94.7 | 0.50 | 0.37 | 9.11 | 0.00 | 0.03 | 0.77 | 10.22 |
| ISSR-4 | 15 | 1 | 16 | 93.8 | 0.50 | 0.37 | 7.44 | 0.00 | 0.03 | 0.79 | 9.56 |
| ISSR-5 | 19 | 1 | 20 | 95.0 | 0.50 | 0.37 | 9.89 | 0.00 | 0.03 | 0.76 | 10.44 |
| ISSR-6 | 16 | 2 | 18 | 88.9 | 0.47 | 0.36 | 6.67 | 0.00 | 0.02 | 0.86 | 7.56 |
| ISSR-8 | 13 | 2 | 15 | 86.7 | 0.41 | 0.33 | 10.67 | 0.00 | 0.03 | 0.50 | 5.56 |
| ISSR-10 | 18 | 3 | 21 | 85.7 | 0.49 | 0.37 | 11.89 | 0.00 | 0.03 | 0.68 | 8.22 |
| ISSR-11 | 16 | 4 | 20 | 80.0 | 0.46 | 0.36 | 12.67 | 0.00 | 0.03 | 0.60 | 7.11 |
| ISSR-12 | 18 | 3 | 21 | 85.7 | 0.49 | 0.37 | 12.00 | 0.00 | 0.03 | 0.67 | 8.00 |
| ISSR-13 | 13 | 2 | 15 | 86.7 | 0.49 | 0.37 | 8.67 | 0.00 | 0.03 | 0.67 | 5.33 |
| ISSR-14 | 10 | 2 | 12 | 83.3 | 0.49 | 0.37 | 6.67 | 0.00 | 0.03 | 0.69 | 4.67 |
| ISSR-18 | 10 | 1 | 11 | 90.9 | 0.50 | 0.37 | 6.00 | 0.01 | 0.03 | 0.71 | 7.33 |
| ISSR-19 | 15 | 2 | 17 | 88.2 | 0.47 | 0.36 | 10.44 | 0.00 | 0.03 | 0.62 | 7.78 |
| ISSR-20 | 17 | 0 | 17 | 100.0 | 0.50 | 0.37 | 7.78 | 0.00 | 0.03 | 0.79 | 9.33 |
| Total | 220 | 32 | 252 | 252 |  |  |  |  |  |  |  |
| Average | 14.66 | 2.13 | 16.8 |  |  |  |  |  |  |  |  |

**Table 7.** Genetic similarities between the nine treatments based on Jaccard’s similarity coefficient based on ISSR primers data. Symbols: C; Control, C15; chlorntraniliprole LC_15_, C50; chlorntraniliprole LC_50_, Cn15; nano-chlorntraniliprole LC_15_, Cn50; Nano-chlorntraniliprole LC_15_, T15; thiocyclam LC_15_, T50; thiocyclam LC_50_, Tn15; nano-thiocyclam LC_15_ and Tn50; nano-thiocyclam LC_50_.

|  | CONTROL | C15 | C50 | T15 | T50 | CN15 | CN50 | TN15 | TN50 |
| --- | --- | --- | --- | --- | --- | --- | --- | --- | --- |
| CONTROL | 100% | -- | -- | -- | -- | -- | -- | -- | -- |
| C15 | 49% | 100% | -- | -- | -- | -- | -- | -- | -- |
| C50 | 56% | 50% | 100% | -- | -- | -- | -- | -- | -- |
| T15 | 54% | 51% | 66% | 100% | -- | -- | -- | -- | -- |
| T50 | 54% | 46% | 65% | 62% | 100% | -- | -- | -- | -- |
| CN15 | 58% | 51% | 65% | 56% | 60% | 100% | -- | -- | -- |
| CN50 | 41% | 47% | 41% | 44% | 42% | 52% | 100% | -- | -- |
| TN15 | 50% | 54% | 51% | 50% | 47% | 55% | 48% | 100% | -- |
| TN50 | 46% | 43% | 52% | 58% | 44% | 47% | 37% | 50% | 100% |

**Fig 6.**
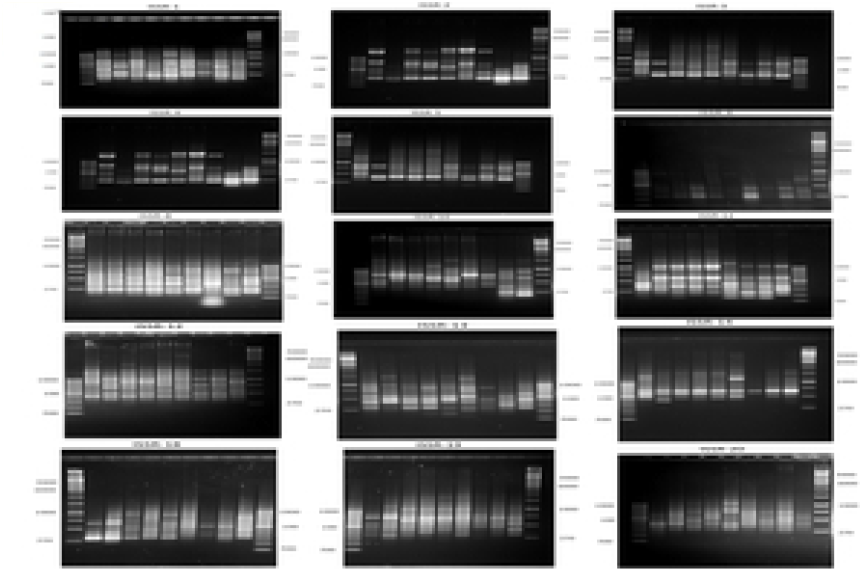
A representative agarose gel where PCR products of the 15 ISSR primers for the nine treatments.

### Analysis of molecular phylogeny

A dendrogram based on the UPGMA cluster analyses of the ISSR data were constructed for the nine treatments (Fig 7). The dendrogram was comprised of two main clusters: the first cluster included only the LC_50_ of chlorantraniliprole, while the second cluster comprised two sub-clusters. The first sub-cluster consisted of only the LC_50_ of nano-thiocyclam, and the second sub-cluster involved two major groups. The first major group included the LC_15_ treatments of chlorantraniliprole, and nano-thiocyclam. The second major group consisted of the control, the LC_15_ treatments of chlorantraniliprole, nano-chlorantraniliprole, and thiocyclam, and the LC_50_ of thiocyclam. Furthermore, the PCA analysis of the ISSR data revealed highly similar results to the cluster analysis. The PCA results indicated that the LC_15_ of nano-chlorantraniliprole and the LC_50_ of thiocyclam were most similar to the control (Fig 8).

**Fig 7.**
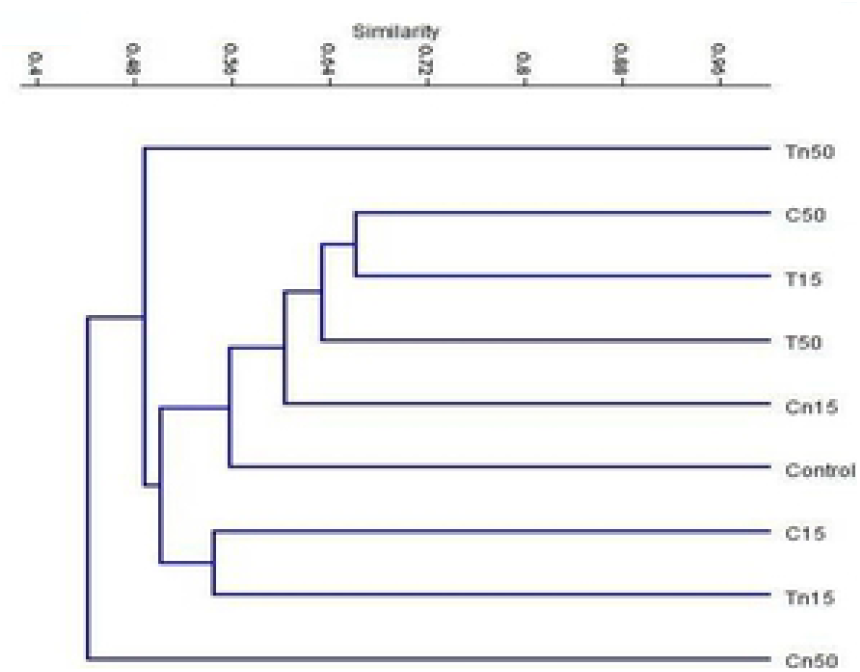
UPGMA cluster analysis based on Jaccard’s similarity coefficient of ISSR analysis of the nine treatments: C; Control, C15; chlorntraniliprole LC_15_, C50; chlorntraniliprole LC_50_, Cn15; nano-chlorntraniliprole LC_15_, Cn50; nano-chlorntraniliprole LC_50_, T15; thiocyclam LC_15_, T50; thiocyclam LC_50_, Tn15; nano-thiocyclam LC_15_ and Tn50; nano-thiocyclam LC_50_.

**Fig 8.**
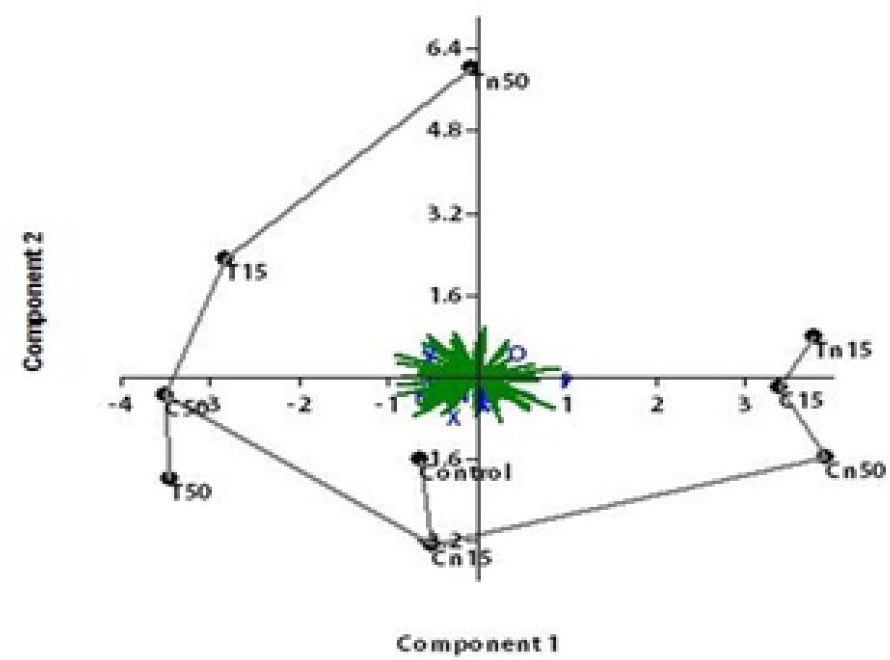
A representative agarose gel where PCR products of the 15 ISSR primers for the nine treatments.

## Discussion

Insecticide efficacy depends on the mode of action, insect species, developmental stage, application methods, and the number of days post treatment [50]. Thus, pesticide nano-formulations may change the nature of chemical pesticides by increasing the efficiency of insect pest control and reducing pesticide application frequency, per the Environmental Protection Agency’s recommendation [51]. This technology is still insufficient and requires a better understanding of its mechanisms prior to field use. We investigated the effects of the lethal and sublethal concentrations of thiocyclam, chlorantraniliprole, and their nano-forms on *A. ipsilon* larval and pupal duration, larvae mortality, adult emergence, reproductive, sex ratio, and oxidative stress enzymes with particular emphasis on DNA mutagenicity.

Chlorantraniliprole (Coragen®) has a novel mode of action [52] that activates insect ryanodine receptors (RyRs), leading to paralysis and mortality in sensitive species [53]. Thiocyclam (Evisect-S®) is a broad-spectrum nereistoxin antagonist that blocks the transmission of cholinergics [52], resulting in insect death. Our results revealed that *A. ipsilon* larvae were more susceptible to the nano-forms than the regular forms for both insecticides (Table 2). This might be due to (1) the nano chlorine within chlorantraniliprole, since chlorine is a non-selective oxidant with a number of effects on the living biota (e.g., reacts with a variety of cellular components, deactivates enzymatic active sites, decreases the biological functions of proteins, and produces deleterious effects on DNA [54]. In some cases, different layers of protein present in insects, larvae or even spores provides protection against chemical attacks, including chlorine, but we speculate that nano-chlorine has a greater effect on the cytoplasmic membrane permeability, causing the loss of refractivity, separating the spore coats from the cortex, extensively discharging Ca+, dipicolinic acid, and DNA, and finally causing lysis to occur, which can lead to cell death and growth inhibition [55].The increased susceptibility to the nano-forms may also be due to (2) the nano-sulfur within thiocyclam, since sulfur destroys an insect’s normal energy-producing bodily functions [56]. Normal pesticides containing sulfur have some caveats, because sulfur can damage plants during hot, dry weather and is incompatible with some other pesticides. In addition, sulfur should not be used on plants that have been sprayed with horticultural oils, as the induced reaction can damage foliage. Sulfur is non-toxic to humans and animals, unless ingested [57]. The nano-form of sulfur does not have the same caveats (at least currently); hence, there is no discrepancy or caveats regarding its application.

Similarly, thiocyclam is a highly effective insecticide against the field strain [58] of *Tuta absoluta* (9.98 mg/L). Likewise, the thiocyclam [59] had the highest efficiency in larvae mortality in field experiments in Iran. In contrast, *A. ipsilon* larvae were more susceptible to chlorantraniliprole and its nano-form than thiocyclam (Table 1). Other lepidopteran pests, including *Helicoverpa armigera, Spodoptera exigua*, and *Spodoptera littoralis* [31,60,61] are also susceptible to this insecticide. The chlorantraniliprole showed harmful activity toward the third instar larvae of *A. ipsilon* [62], whereas its LC_50_ was 0.187 μg/g 72 h post treatment. Generally, pest susceptibility to chemical insecticides is affected by various factors, including nutrition type, size, and physiological status of the host [63].

Sublethal effects of insecticides could be considered common toxicological phenomena and represent a secondary effect of insecticide application [62,64]. This phenomenon may occur in insects due to exposure to degraded insecticide after itsinitial application on crops [31]. When second instar larvae of *A. ipsilon* are exposed to sublethal concentrations of chlorantraniliprole, thiocyclam, and their nano-forms, the developmental rates of the larvae and pupae were substantially extended (Table 2) in the original form of both insecticides compared to their nano-forms. These results agree with [62], who found that the development of *A. ipsilon* larvae was prolonged by low concentrations of chlorantraniliprole. Chlorantraniliprole has also been shown to prolong the development times of some lepidopteran insect pests, including; *Plutella xylostella, S. littoralis*, and *Spodoptera cosmioides* [65,66,67]. In this study, both insecticides and their nano-forms did not show any significant differences in other biological parameters, including pupal weight, pupation%, and emergence%, except for the LC_50_ of nano-thiocyclam, which exhibits a significant difference in pupation% and emergence%. Additionally, the nano-form of both insecticides at LC_50_ showed a significant difference in the number of eggs laid per female and fertility. Several studies have also shown that low concentrations of insecticides may affect the reproductive activity of insects (e.g., *Helicoverpa assulta, Mamestra brassica*, and *S. littoralis* [31,68,33].

In the last decade, pesticide-induced oxidative stress has been a focus of toxicological research as a possible toxicity mechanism that causes a final manifestation of a pro-oxidant and antioxidant defense mechanism imbalance [69]. Pesticide intoxication causes a derangement of different antioxidant mechanisms in various tissues [69]; however, exposure to sublethal insecticide concentrations can induce oxidative stress enzymes to increase (e.g., SOD, CAT, and GR) [70]. Our results indicate that the CAT activity was significantly increased in both the insecticides and their nano-forms. Significantly increased SOD activity was observed in the LC_50_ of both insecticides and their nano-forms. Also, significantly higher GR activities were observed in nano-chlorantraniliprole and nano-thiocyclam. The antioxidant enzyme systems, such as CAT or SOD, play an essential role in detoxifying harmful agents to counter the damaging effects of ROS [71].

Similar to the elevated antioxidant levels, lipid peroxide activity was significantly increased in MDA, which is the main oxidation result from the peroxidation of unsaturated fatty acids. This represents the oxidative effect on different organisms [72,73] in the LC_50_ of both insecticides and their nano-forms. We observed no significant differences between the insecticides and their nano-forms at the lowest tested concentration (LC_15_), which reflects the protective effects of antioxidants in *A. ipsilon* larvae. The antioxidant system can reduce MDA levels, which contributes to the accumulation of active oxygen and inhibits antioxidase activity [74]. The accumulation of MDA can be reduced by antioxidant enzymes, such as SOD and CAT. Other studies have demonstrated that toxic oxygen metabolism can mediate lipid peroxidation [75].

The ISSR-PCR technique was used as an effective marker to investigate the genetic mutagenicity levels between the second instar larvae subjected to different insecticidal treatments (chlorantraniliprole, thiocyclam and their nano-forms) compared to the control (untreated). One of the benefits of ISSR is that it performs a qualitative evaluation of DNA variability through the differences in amplification profiles [44]. Many studies reported that the exposure of insects to particular insecticides at different concentration levels might lead to damages or changes in genomic DNA sequences (e.g., insertions, deletions, substitutions, or rearrangements), resulting in changes in the ISSR profile (e.g., the presence or absence of certain bands or variations of the band intensity) [25]. In addition, the possible occurrence of point mutations at the oligonucleotide annealing site may cause the absence of bands due to the loss of a priming site [76]. In this study, 15 ISSR primers were used to detect genetic mutagenicity levels and screen the degree of polymorphism between treated and untreated larvae. Furthermore, to establish the relationship between the insecticides, the ISSR study results were used to create a dendrogram. The results showed that the lowest mutagenic insecticidal effects on the insect DNA was observed in nano-the LC_15_ of chlorantraniliprole compared to the control. In contrast, the most aggressive (highest) mutagenic effect was observed in the LC_50_ of nano-chlorantraniliprole, followed by the LC_50_ of nano-thiocyclam LC_50_ compared to the control. It was elucidated that the PCA analysis of ISSR data revealed a consistent result to the obtained by the dendrogram topology. Moreover, it was found that the LC_15_ of nano-chlorantraniliprole and the LC_50_ of thiocyclam exhibited almost the same mutagenic effects as the control.

It is expected that the polymorphic differences observed in the ISSR patterns may be attributed to changes in primer binding sites or DNA structures, or to DNA damage caused by insecticide exposure. In addition, it could also be due to the blocking of DNA replication, the presence of a large number of chromosomal lesions, such as large rearrangements (e.g., deletion, inversion, or translocation), or un-repaired DNA damage due to direct exposure to different insecticidal treatments [77]. The ISSR studies demonstrated their ability and effectiveness to detect DNA damage and changes caused by the studied insecticides.

## Conclusion

In summary, we developed insecticide nanometerization to improve the biological activity of conventional insecticides. Our obtained results represent a promising step toward developing safe and efficient nanoinsecticides. Further investigations are still needed to determine their effects on the environment. To the best of our knowledge, this research is a pioneer case-study that examined and analyzed the genome-wide DNA mutability, biochemical effects, and toxicity levels of chlorantraniliprole, thiocyclam, and their nano-forms for the control of *A. ipsilon*.

## Acknowledgments

Authors are thankful for the Egyptian Knowledge Bank (EKB) for improving the manuscript considerably, including English language and grammar.

